# Uracil-DNA Glycosylase of Murine Gammaherpesvirus 68 Binds Cognate Viral Replication Factors Independently of its Catalytic Residues

**DOI:** 10.1101/2023.05.19.541466

**Authors:** Kyle R. Smith, Somnath Paul, Qiwen Dong, Orchi Anannya, Darby G. Oldenburg, J. Craig Forrest, Kevin M. McBride, Laurie T. Krug

**Affiliations:** HIV and AIDS Malignancy Branch, National Cancer Institute, Bethesda, MD, USA; Department of Microbiology & Immunology, Stony Brook University, Stony Brook, NY, USA; Department of Epigenetics and Molecular Carcinogenesis, The University of Texas MD Anderson Cancer Center, Houston, Texas, USA; Molecular and Cellular Biology Program, Stony Brook University, Stony Brook, NY, USA; Department of Physiology and Biophysics, Molecular and Cellular Biology Program, Stony Brook University, Stony Brook, NY, USA; Gundersen Medical Foundation, Gunderson Health System, LaCrosse, Wisconsin, USA; Department of Microbiology & Immunology, University of Arkansas for Medical Sciences, Little Rock, Arkansas, USA

**Keywords:** Gammaherpesvirus, lytic replication, uracil-DNA glycosylase, viral DNA polymerase, viral DNA polymerase processivity factor, viral replication compartment, virus-host interactions

## Abstract

Herpesviruses are large double-stranded DNA viruses that encode core replication proteins and accessory factors involved in nucleotide metabolism and DNA repair. Mammalian Uracil-DNA glycosylases (UNG) excise deleterious uracil residues from their genomic DNA. Each herpesvirus UNG studied to date has demonstrated conservation of the enzymatic function to excise uracil residues from DNA. We previously reported that a murine gammaherpesvirus (MHV68) with a stop codon in *ORF46* (ORF46.stop) that encodes for vUNG was defective in lytic replication and latency *in vivo.* However, a mutant virus that expressed a catalytically inactive vUNG (ORF46.CM) had no replication defect, unless coupled with additional mutations in the catalytic motif of the viral dUTPase (ORF54.CM). The disparate phenotypes observed in the vUNG mutants led us to explore the non-enzymatic properties of vUNG. Immunoprecipitation of vUNG followed by mass spectrometry in MHV68-infected fibroblasts identified a complex comprised of the cognate viral DNA polymerase, vPOL encoded by *ORF9*, and the viral DNA polymerase processivity factor, vPPF encoded by *ORF59*. MHV68 vUNG colocalized with vPOL and vPPF in subnuclear structures consistent with viral replication compartments. In reciprocal co-immunoprecipitations, the vUNG formed a complex with the vPOL and vPPF upon transfection with either factor alone, or in combination. Last, we determined that key catalytic residues of vUNG are not required for interactions with vPOL and vPPF upon transfection or in the context of infection. We conclude that the vUNG of MHV68 associates with vPOL and vPPF independently of its catalytic activity.

**IMPORTANCE:** Gammaherpesviruses encode a uracil-DNA glycosylase (vUNG) that is presumed to excise uracil residues from viral genomes. We previously identified the vUNG enzymatic activity, but not the protein itself, as dispensable for gammaherpesvirus replication *in vivo*. In this study, we report a non-enzymatic role for the viral UNG of a murine gammaherpesvirus to form a complex with two key components of the viral DNA replication machinery. Understanding the role of the vUNG in this viral DNA replication complex may inform the development of antiviral drugs that combat gammaherpesvirus associated cancers.

## INTRODUCTION

Herpesviruses are large, double-stranded DNA viruses that encode proteins necessary for viral DNA replication (1). The viral DNA polymerase (vPOL), DNA polymerase processivity factor (vPPF), single-strand DNA binding protein (vSSBP), and a helicase-primase complex comprise the core replication machinery. Herpesviruses also encode accessory factors that are homologs of host enzymes involved in nucleotide metabolism, including a thymidine kinase (vTK), ribonucleotide reductase large and small subunits (vRNR-L and vRNR-S), a deoxyuridine 5′-triphosphate nucleotidohydrolase (vDUT), and a DNA repair protein, the uracil-DNA Glycosylase (vUNG). While herpesvirus accessory factors are generally dispensable for replication in rapidly dividing cells, they promote replication in more restrictive conditions such as low multiplicity infections and non-dividing cells that are common in the host (1-5).

Uracilated DNA arises from misincorporation, spontaneous cytosine deamination, or actions by the host enzymes activation-induced cytidine deaminase (AID) in germinal center B cells and members of the apolipoprotein B mRNA-editing enzyme catalytic polypeptide (APOBEC) family. Left unrepaired, uracils in DNA drive mutations, lesions, and genomic instability (6-8). APOBEC cytidine deaminases act on herpesvirus genomes to induce mutations and impair replication (9-13). Eurakyotes encode host UNGs to surveil and excise uracils from single-stranded DNA and U:A or U:G mismatches in double-stranded DNA to initiate repair of the abasic site using the base excision repair (BER) machinery (14-16). Herpesviruses encode UNG homologs with conserved enzymatic domains and overlapping biochemical properties. Uracil excision by HSV-1 vUNG (*UL2*) is followed by repair in coordination with vPOL (*UL30*) and host BER machinery, including AP endonuclease and the ligase IIIα-XRCC1 complex (17). Disruption of the catalytic domain of HSV-1 vUNG increases instability in the viral genome following serial passage in cell culture (3). The vUNG of murine gammaherpesvirus 68 (MHV68) excises uracil and functionally substitutes for host UDG following AID deaminase conversion of cytosine to uracil to trigger immunoglobulin isotype class-switch recombination in B cells (18).

The vUNG DNA repair protein is largely dispensable for replication in rapidly proliferating cells but is required for replication in resting cells or restrictive tissues *in vivo*. The varicella-zoster virus (VZV) UNG (*ORF59*) is dispensable for replication in human melanoma cells and monkey kidney fibroblast cells and a UNG inhibitor does not impact VZV replication (19, 20). Similarly, HSV-1 vUNG is dispensable for replication in immortalized murine fibroblasts and hamster kidney cells (3, 21, 22). However, an HSV-1 UNG mutant displayed decreased replication and neurovirulence in mice (22). A human cytomegalovirus mutant with a deletion in *UL114* that encodes its cognate vUNG exhibitd delayed kinetics in viral DNA synthesis and virion production in primary human embryonic cells and in serum-starved primary human foreskin fibroblasts (5, 23, 24). Likewise, the MHV68 vUNG is not essential for replication in cell culture but is required for pathogenesis *in vivo* (18, 25).

We previously reported that a recombinant MHV68 lacking vUNG due to a stop codon engineered in *ORF46* (ORF46.stop) exhibits a dose-dependent lytic replication defect in the lungs and a route-dependent defect in latency establishment in spleens of mice following intranasal infection (18). To determine if the enzymatic activity of vUNG was responsible for the replication defect of the ORF46.stop virus, we introduced catalytic mutations into two enzymatic domains of vUNG to generate a recombinant vUNG catalytic mutant (ORF46.CM) virus (25). The catalytic mutations, D85N in the water activating loop responsible for hydrolytic displacement of the uracil residue and H207L in the DNA binding domain, eliminated uracil excision *in vitro* (25). Unexpectedly, the ORF46.CM mutant did not reproduce the replication defect of the ORF46.stop virus in cell culture or in the lungs of infected mice.

The failure of the ORF46.CM mutant to recapitulate the replication defect of the ORF46.stop mutant led us to explore non-enzymatic functions of MHV68 vUNG. The vUNGs of HSV-1, HCMV, and EBV interact and colocalize with their viral replication factors, vPOL and vPPF (23, 25-28). We hypothesized that the vUNG of MHV68 would interact with cognate viral replication factors and, further, that these interactions would be independent of its catalytic activity. Here, we report an unbiased immunoprecipitation mass spectrometry (IP-MS) approach that identified vPOL and vPPF as viral factors that interact with the MHV68 vUNG. We used recombinant viruses with GFP tagged vPOL and vPPF to examine colocalization with vUNG following infection. We also investigated the interaction of vUNG with vPOL or vPPF in the absence of other viral replication factors. Finally, we assessed the requirement of the catalytic activity of vUNG for co-localization and complex formation with these cognate factors. In sum, we report a non-enzymatic function of the vUNG DNA repair protein to complex with the vPOL and vPPF of MHV68.

## RESULTS

### Identification of vUNG binding partners by IP-mass spectrometry

MHV68 vUNG-protein interactions that occur during lytic replication have not been defined. To identify vUNG binding partners in an unbiased manner, co-immunoprecipitation followed by mass spectrometry (co-IP/MS) was performed on *de novo* infected fibroblasts. Flow cytometry with a monoclonal antibody (vUNG-C1 mAb) in conjunction with a YFP reporter was used to identify the timepoint at which vUNG expression levels were highest in infected cells. The expression of vUNG and infected cells increased with time (Fig. 1A-B) and cell viability was comparable between mock and infected cells up to 36 hours post-infection (hpi) (Fig. 1C). Immunoprecipitation of protein lysates from MHV68 infected fibroblasts at 36 hpi with the vUNG-C1 mAb (29) followed by immunoblot with polyclonal vUNG anti-serum detected the expected 28 kDa protein for vUNG (Fig. 1D). These 36 hpi IP samples were next subjected to MS analysis. For each protein the false discovery rate was set to 0.1% and at least ten unique peptide hits were required. vPOL and vPPF had the most abundant peptide spectra in terms of number of unique peptides and peptide coverage. vPOL and vPPF were the only viral proteins enriched two-fold (Log_2_FC=1.0) over IgG control (Fig. 1E). These results suggest that vPOL and vPPF engage vUNG during MHV68 *de novo* lytic infection.

**FIG 1.**
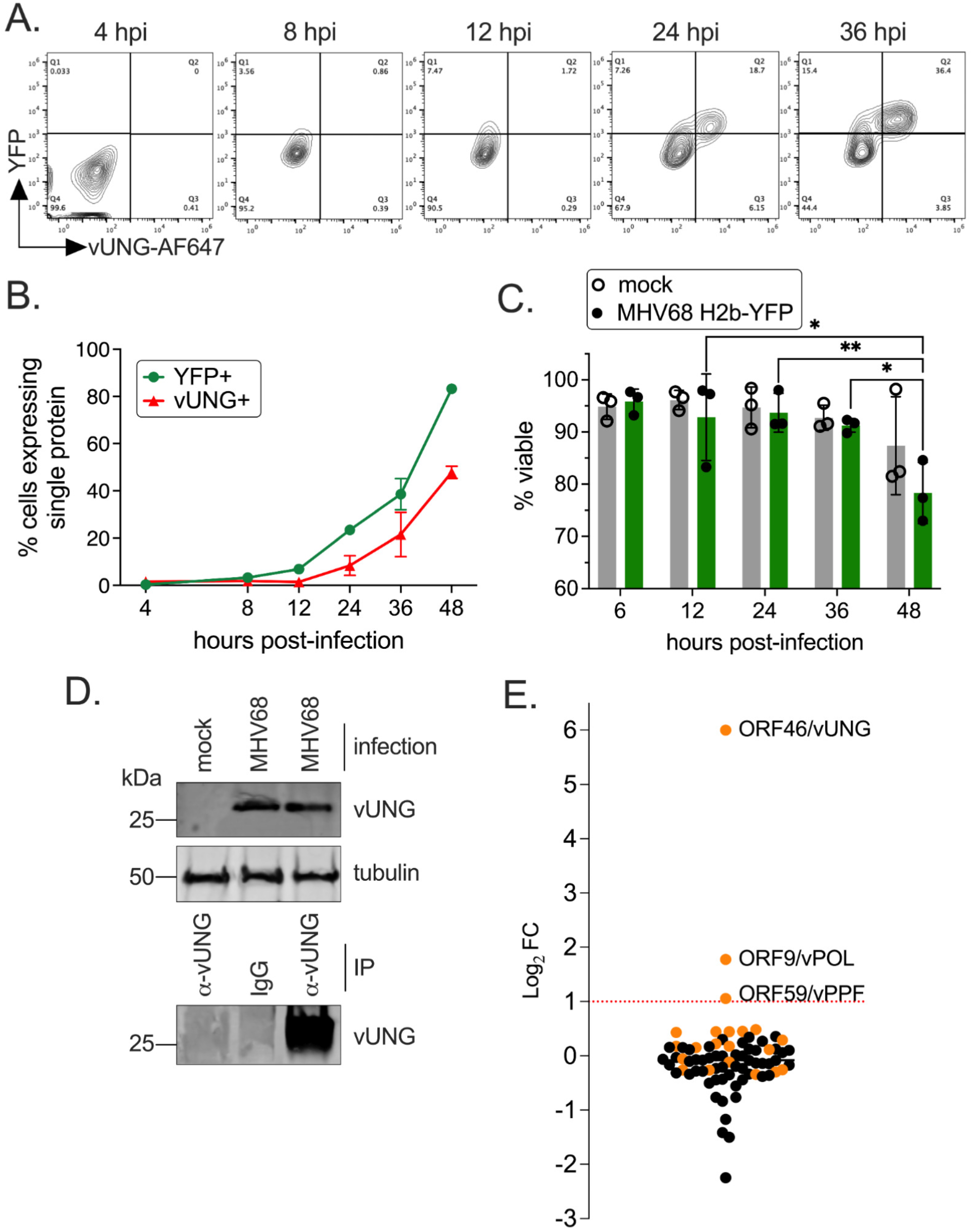
The MHV68 vUNG interacts with the vPOL and vPPF upon *de novo* lytic infection. NIH 3T12 cells were infected with MHV68 H2b-YFP at an MOI of 3.0. (A) Cells were fixed and stained for flow cytometry at the indicated hpi. Plots indicate the gating strategy used to identify YFP+ infected cells on the y-axis and cells expressing vUNG detected by the vUNG-C1 mAb conjugated to AF647+ on the x-axis. (B) Line graph of percentage of cells with YFP, or vUNG from flow data collected in (A). (C) Cell viability of mock and infected cells was measured by exclusion of propidium iodide by flow cytometry. Symbols represent three biological replicates +/- SEM. Statistical significance was evaluated by two-way Anova with Tukey’s multiple comparisons test. *p<0.05, **p<0.01 (D) Immunoprecipitation of the vUNG using the vUNG-C1 mAb from infected NIH 3T12 cells at 36 hpi. (E) Enriched peptides identified by mass spectrometry with a log_2_ fold change following IP of vUNG compared to control IgG. The orange spheres denote MHV68 specific peptides detected, while the black spheres denote host/murine peptides.

### vUNG complexes with cognate viral DNA replication factors vPOL and vPPF

To investigate whether vUNG colocalizes with vPOL and vPPF during MHV68 infection, recombinant viruses were generated to express vPOL or vPPF as C-terminal GFP fusion proteins. Infectious particle production and viral protein expression were compared following infection of NIH 3T12 fibroblasts with either WT or recombinant MHV68 vPOL-GFP and vPPF-GFP viruses. Viral replication kinetics were similar between the viruses expressing vPOL-GFP or vPPF-GFP compared to WT virus, with minor deficits for vPOL-GFP virus at 12 and 36 hpi (Fig. 2A). Both vPOL-GFP and vPPF-GFP migrated at the expected relative molecular weights based on predicted size and the addition of GFP. The expression of vUNG and other lytic antigens was similar between the WT and GFP tagged viruses at both 24 and 36 hpi (Fig. 2B).

**FIG 2.**
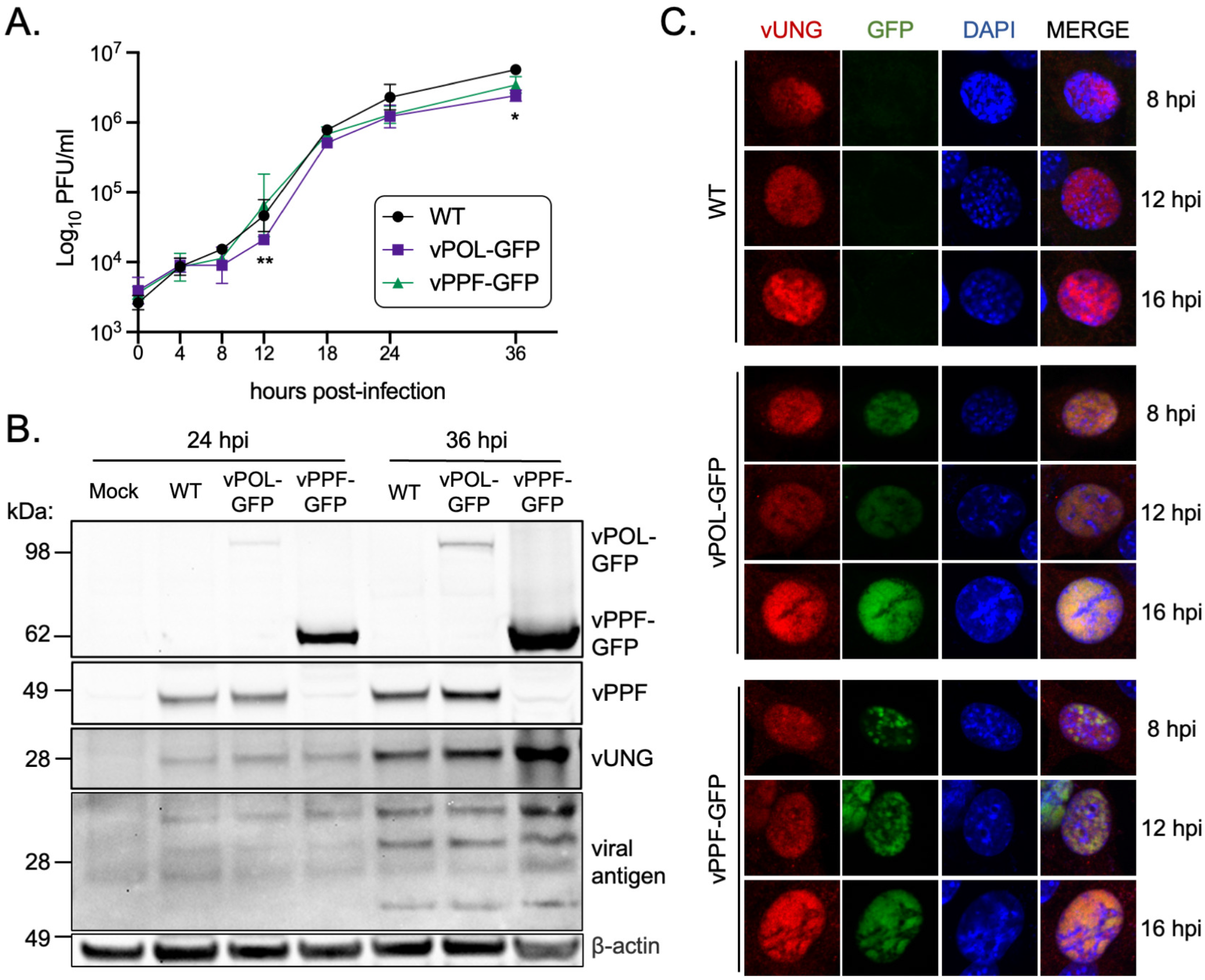
Characterization and immunofluorescence imaging of recombinant MHV68 expressing GFP tagged vPOL and vPPF. (A) Single-step growth curve to compare replication of wild-type MHV68 (WT) or recombinant MHV68 expressing GFP-fusions to vPOL (viral DNA polymerase encoded by *ORF9*) or vPPF (viral DNA polymerase processivity factor encoded by *ORF59*) in NIH 3T12 fibroblasts, MOI 3.0. Symbols represent three biological replicates +/- SEM. Statistical significance was evaluated by two-way Anova with Tukey’s multiple comparisons test and differences were noted between ORF9-GFP and WT at 12 and 36 hpi, and between ORF9-GFP and ORF59- GFP at 12 hpi; *, p<0.05; **p<0.01 (B) Timecourse of viral protein expression by immunoblot under conditions described in A. (C) Immunofluorescence timecourse of viral protein colocalization in NIH 3T3 cells infected with the indicated viruses, MOI 3.0. vUNG was detected with vUNG pAb followed by secondary AF568; vPOL and vPPF fusion proteins were detected via GFP expression without antibodies; DNA was stained with DAPI.

The recombinant MHV68 GFP fusion viruses were used to investigate vPOL and vPPF colocalization with the vUNG over a timecourse of lytic infection by confocal microscopy. The essential viral replication factors vPOL and vPPF are well-characterized markers of virus replication compartments (vRCs) in herpesvirus infections. The size and contiguity of the nuclear structures composed of vPOL-GFP and vPPF-GFP increased with time, leading to large globular structures demarcated from DAPI-stained chromosomal DNA by 16 hpi (Fig. 2C). vUNG expression increased with time and overlapped spatially with both vPOL-GFP and vPPF-GFP at each timepoint. These findings indicate that vUNG colocalizes with vPOL and vPPF in vRC-like structures during lytic MHV68 infection.

Having observed localization of vUNG with vPOL and vPPF, we tested whether viral DNA synthesis was required for colocalization. Infected NIH 3T3 fibroblasts were treated with phosphonoacetic acid (PAA), a specific inhibitor of viral DNA synthesis, immediately after inoculation to block viral DNA replication (Fig. 3A-B, top three rows). In the presence of PAA, vUNG, vPOL and vPPF were largely confined to small puncta at 9 hpi that did not expand with time; vUNG localization largely overlapped with vPOL and vPPF. This contrasted with untreated cells (Fig. 3A-B, bottom three rows) where vUNG and vPOL, and vUNG and vPPF colocalize in vRCs that increased in size along with the nuclear abundance of viral proteins from early to late hpi as previously observed (30, 31). Representative line graphs following Nikon NIS-Elements analysis indicate strong overlap in intensity and distance for each viral factor at 17 hpi (Fig. 3A-B, bottom panels).

**FIG 3.**
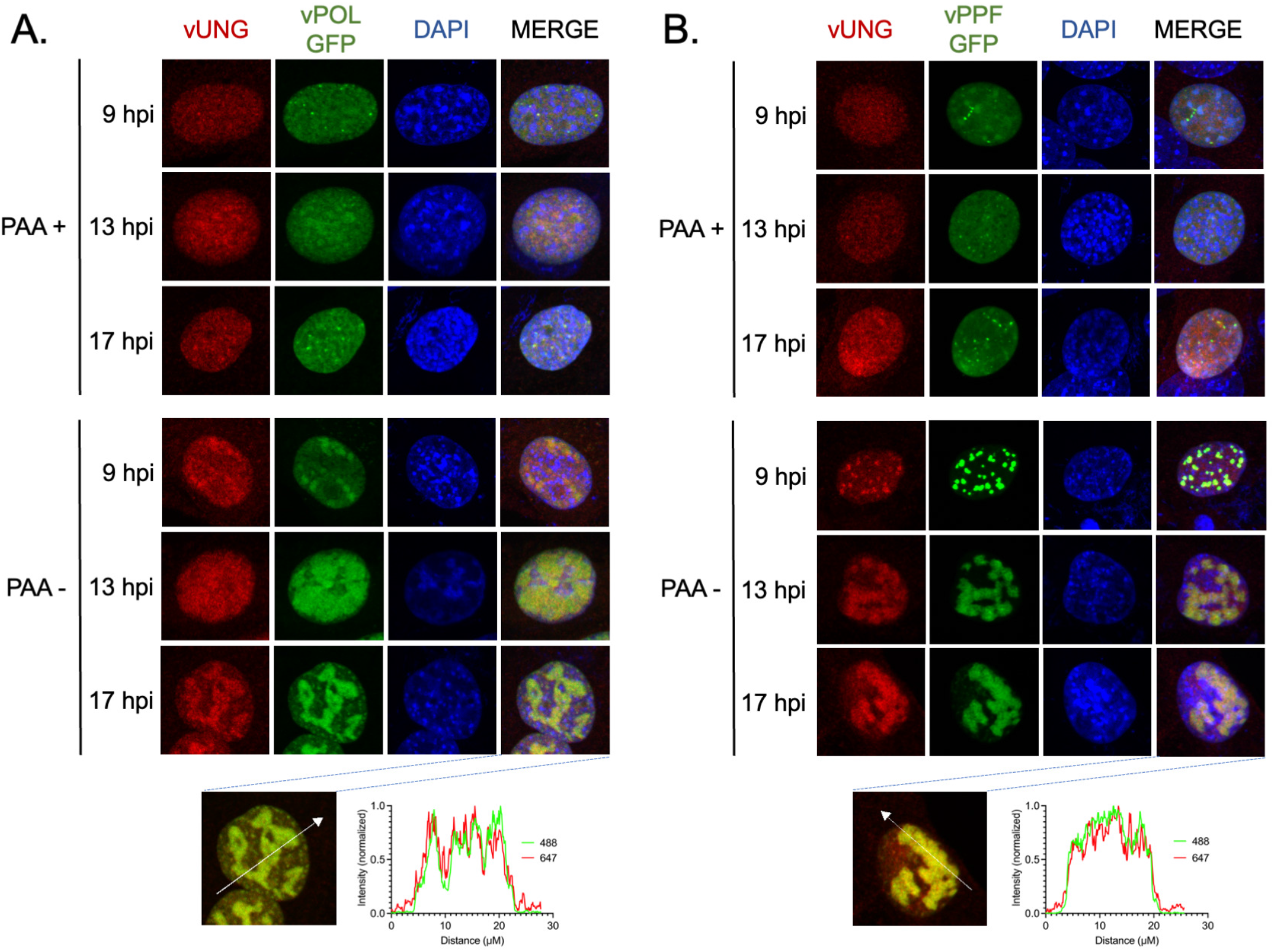
vUNG colocalizes with the viral DNA polymerase and viral DNA polymerase processivity factor in infected fibroblasts. (A-B) NIH 3T3 cells were infected with recombinant MHV68 virus expressing vPOL-GFP (A) or vPPF-GFP (B) at an MOI of 3.0, and treated with 300 ug/mL PAA at 1 hpi where indicated. Cells were fixed and stained for immunofluorescence imaging at the indicated hpi. vUNG was detected with vUNG-C1 mAb conjugated to AF647; vPOL and vPPF fusion proteins were detected via GFP expression without antibodies; DNA was stained with DAPI. Nikon NIS-Elements line analysis of the indicated cells at 17 hpi are located below each panel (A-B).

### vUNG interacts with vPOL and vPPF independent of other viral factors

To assess whether the interaction of vUNG with vPOL and vPPF requires additional viral factors, co-IP was performed upon transient expression in HEK-293T cells. Immunoprecipitation of vUNG co-precipitated FLAG-vPOL and vPPF-V5 (Fig. 4A). A reciprocal co-IP of vPPF-V5 with V5 antibodies captured FLAG-vPOL and vUNG-MYC (Fig. 4B). We next interrogated the ability of vUNG to complex with either vPOL or vPPF upon co-transfection of paired expression constructs. As expected, vPOL and vPPF formed a complex in reciprocal IP experiments; vPPF-V5 and FLAG-vPOL were pulled-down with FLAG mAb (Fig. 4C) or with the V5 mAb (Fig. 4D). Immunoprecipitation of vUNG co-precipitated FLAG-vPOL and vUNG-MYC (Fig. 4E), and IP of vPPF co-precipitated FLAG-vUNG and V5-vPPF (Fig. 4F). These data indicate that vUNG, vPOL and vPPF interact in pairs as well as in a complex independent of active viral DNA replication and other viral proteins.

**FIG 4.**
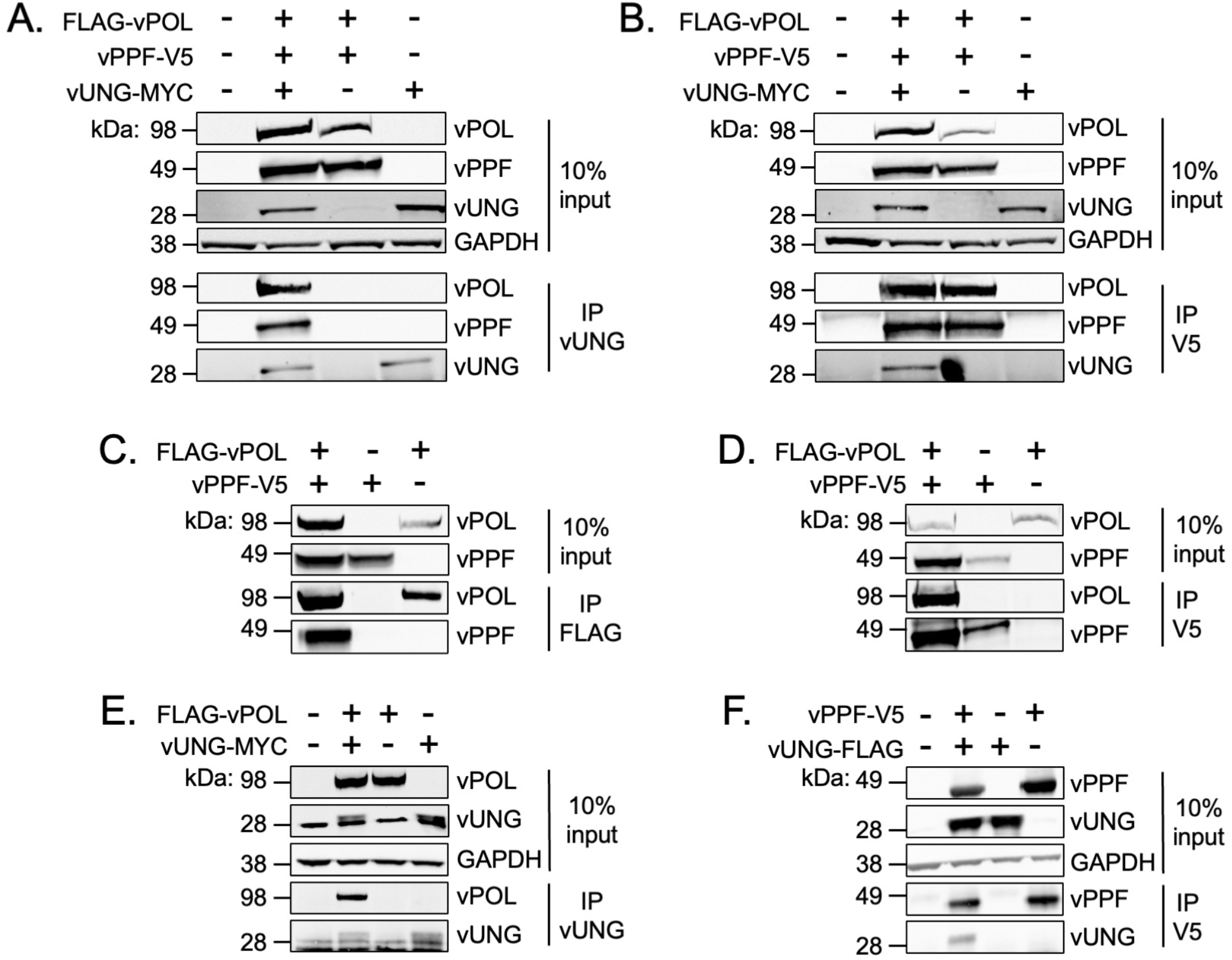
vUNG interacts in complex with vPOL and vPPF. At 48 h post-transfection, protein lysates were DNAse treated and sonicated prior to immunoprecipitation (IP) and immunoblot analysis. (A) IP with anti-vUNG C1 mAb or anti-V5 mAb (B) in lysates from co-transfections with FLAG-vPOL, vPPF-V5, and vUNG-MYC. (C & D) IP with anti-FLAG mAb (C) or anti-V5 mAb (D) in lysates from co-transfections with FLAG-vPOL and vPPF-V5. (E) IP with anti-FLAG mAb in lysates from co-transfections with vUNG-MYC and FLAG-vPOL. (F) IP with anti-V5 mAb in lysates from co-transfections with vUNG-FLAG and vPPF-V5.

### The complex of vUNG, vPOL and vPPF forms independently of the enzymatic activity of vUNG

Given that the vUNG interacted with the viral DNA replication factors vPOL and vPPF in the absence of ongoing viral DNA synthesis, we next tested if these interactions would occur independently of the enzymatic activity of vUNG. We performed immunofluorescence imaging following infection with either the MHV68 ORF46 marker rescue virus (ORF46.MR) expressing WT vUNG (Fig. 5A) or the MHV68 ORF46.CM virus expressing the catalytic mutant (CM) form of vUNG (Fig. 5B). Both WT vUNG and vUNG.CM colocalized with vPPF from 9-17 hpi (Fig. 5A-B). Colocalization with vPPF in the context of *de novo* fibroblast infection is independent of vUNG catalytic activity.

**FIG 5.**
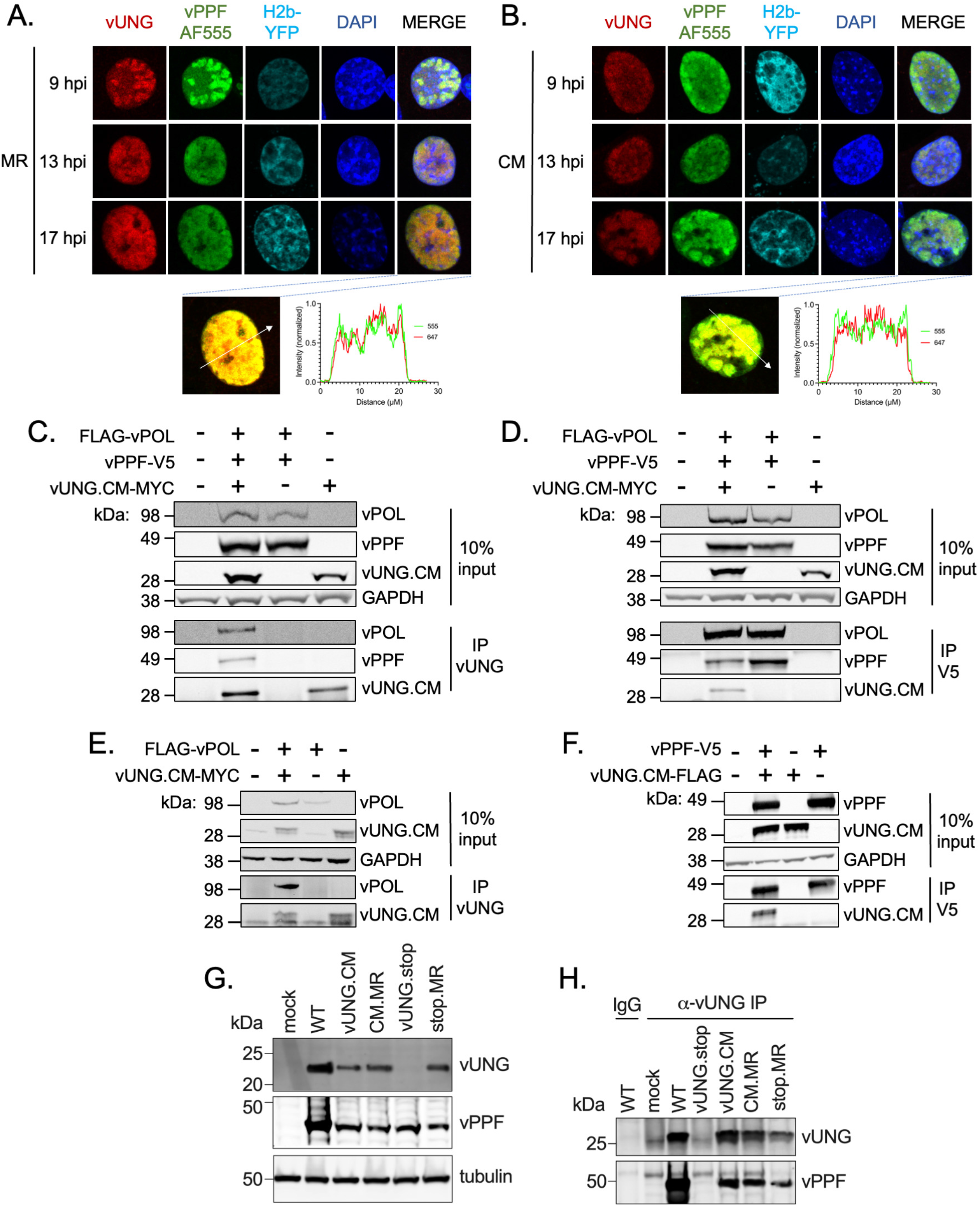
vUNG complexes with vPOL and vPPF independent of vUNG enzymatic activity. (A-B) NIH 3T3 cells were infected with recombinant MHV68 ORF46CM.MR virus expressing a wild-type vUNG (A) or with MHV68 ORF46.CM expressing the vUNG catalytic mutant (CM) (B) at an MOI of 3.0. vUNG was detected with vUNG-AF647 mAb; vPPF was detected with the vPPF pAb followed by secondary AF555; DNA was stained with DAPI. Nikon NIS-Elements line analysis of the indicated cells at 17 hpi are located below each panel (A-B). (C-F) HEK-293T cells were transfected with the indicated expression constructs. At 48 h post-transfection, protein lysates were DNAse treated and sonicated prior to immunoprecipitation and immunoblot analysis. IP with anti-vUNG C1 mAb (C) or anti-V5 mAb (D) in lysates from co-transfections with FLAG-vPOL, vPPF-V5, and vUNG.CM-MYC. (E) IP with anti-vUNG mAb in lysates from co-transfections with vUNG.CM-MYC and FLAG-vPOL. (F) IP with anti-V5 mAb in lysates from co-transfections with vUNG.CM-FLAG and vPPF-V5. (G) NIH 3T12 cells were infected with the indicated viruses for 36 h and protein lysates were subjected to SDS-PAGE. Tubulin was used as a loading control. (H) Co-IP of NIH 3T12 infected cell lysates at 36 hpi with anti-vUNG-C1 or IgG control. vUNG was detected with the vUNG pAb, and vPPF was detected with the anti-vPPF pAb. Data represents three biological replicates.

We next used transient transfection in HEK-293T cells to determine if complex formation with vPOL in addition to vPPF occurs independent of the enzymatic activity of vUNG. Co-IPs performed with the vUNG.CM precipitated FLAG-vPOL and vPPF-V5 (Fig. 5C), and reciprocal IP of vPPF-V5 likewise captured both vUNG.CM-MYC and FLAG-vPOL (Fig. 5D). IP of lysates following co-transfection of vUNG.CM-MYC and FLAG-vPOL with vUNG-C1 mAb led to pull-down of FLAG-vPOL with vUNG.CM (Fig. 5E). An interaction between vUNG.CM and vPPF-V5 was observed following IP of vUNG.CM with anti-V5 (Fig. 5F). Taken together, the vUNG, vPOL and vPPF interact in pairs and in a complex independent of the enzymatic activity of vUNG, active viral DNA replication, and other viral proteins.

The observation that the vUNG.CM colocalizes with vPPF (Fig. 5B) and interacts with vPPF upon transfection (Fig. 5C-D, Fig. 5F) led us to test for this interaction in the context of infection. The expression of vUNG and vPPF was comparable following infection with WT, CM.MR or the vUNG.CM (Fig. 5G). The vUNG was not detected upon infection with the MHV68 recombinant with a stop codon in ORF46 (vUNG.stop) but detection was restored upon infection with the vUNG.stop marker rescue revertant (stop.MR). IP with the vUNG-C1 mAb was used to examine complex formation with untagged WT or the untagged catalytic mutant form of vUNG upon infection with WT, MR, or vUNG.CM. Comparable levels of vPPF were detected upon pull-down of either WT or catalytic mutant vUNG upon infection with CM.MR or vUNG.CM viruses (Fig. 5H). IP with the vUNG-C1 mAb led to no pull-down of vUNG or vPPF upon infection with vUNG.stop, in contrast to restoration of the complex upon infection with stop.MR (Fig. 5H). Together these data confirm the interaction of vUNG.CM and vPPF in the context of infection, irrespective of the catalytic activity of the vUNG.

## DISCUSSION

In the current study, we identified non-enzymatic functions of the vUNG encoded by the model gammaherpesvirus pathogen MHV68. We report that vUNG readily complexes with vPOL and vPPF in the context of infection, independently of key catalytic residues. This indicates that the vUNG may provide scaffolding functions for the viral DNA replication machinery. Our data offers one potential explanation for the failure of the ORF46.CM mutant to phenocopy the severe replication defect of the ORF46.stop mutant *in vivo* (25).

Through an unbiased co-IP/MS approach and comprehensive validation, we demonstrated that vUNG binds each of the viral DNA replication factors vPOL and vPPF, irrespective of active viral DNA synthesis. These data are consistent with findings for vUNGs of HSV-1 (*UL2*), HCMV *(UL114)*, and EBV *(BKRF3)* (23, 26-28, 31). HSV-1 vUNG co-IPs with vPOL (*UL30*) and vPPF (*UL42)* upon transfection of tagged vUNG followed by HSV-1 infection of HEK-293 cells (27). Infection of human foreskin fibroblasts with recombinant HCMV expressing FLAG tagged vUNG followed by co-IP with anti-FLAG Ab led to pull-down of both vPPF (*UL44*) and vPOL (*UL54*) (28). Paired interactions also were observed *in vitro* with HCMV vUNG and vPPF or vPOL as purified recombinant proteins; the interaction of vUNG with vPPF required the addition of DNA (28, 31). For EBV, epitope-tagged vUNG interacts with vPPF (*BMRF1)* or vPOL *(BALF5)* following transfection in HEK-293Ts and with vPOL following reactivation (26). Further, the interaction between vUNG and vPPF occurred irrespective of mutations in the vUNG enzymatic domain. We extend these findings by demonstrating the formation of a complex containing MHV68 vUNG and both the vPOL and vPPF following transfection conditions that are devoid of viral DNA or other viral factors. The interaction of vUNG with vPOL and vPPF occurs in the context of transfection with enzymatically inactive vUNG.CM. Furthermore, we confirmed that the interaction of non-tagged endogenous levels of vUNG with vPPF occurs independent of enzymatic activity upon *de novo* productive infection.

Our finding that the MHV68 vUNG colocalizes with vPOL and vPPF is consistent with reports for the colocalization of the vUNG of HSV-1, HCMV and EBV with their respective vPOL and vPPFs. HSV-1 vUNG colocalized with vPOL following transfection of Vero cells with vUNG, vPOL and the six other core HSV-1 replication factors (27). HCMV vUNG localized to vRCs in the nucleus with a similar pattern of distribution as vPPF (23). These vRCs expanded from early nuclear puncta to large globular nuclear domains (31). This morphology was reflective of our findings of puncta at early timepoints and with infection in the presence of PAA that expanded to fill large portions of the nucleus by late timepoints. Nuclear localization of the EBV vUNG with either vPOL or vPPF was reported following plasmid transfection in HeLa cells. WT EBV vUNG or a vUNG with Q90L/D91N mutations in the enzymatic domain colocalized with vPOL upon reactivation from a nasopharyngeal carcinoma cell line (26). We extend these findings by reporting that the MHV68 vUNG colocalizes with the vPPF in the context of *de novo* infection. Similar to the EBV studies, we find this colocalization does not require residues D85 and H207 that are critical for the enzymatic activity of MHV68 vUNG (25).

HSV-1 vUNG was observed to localize to nascent viral DNA during productive infection (32). vUNG also coordinates BER in part via interaction with its cognate viral DNA polymerase and host factors *in vitro* (17). These data are consistent with a model whereby vUNG excision of uracil residues triggers BER to prevent recruitment of error-prone translesion DNA polymerases that would otherwise place the genomes at risk for mutation (9, 12). In the case of HCMV infection, the Y-family of translesion DNA polymerases are largely deleterious to viral DNA synthesis while the polymerase ζ complex stabilizes viral genome integrity (33). Accelerated native isolation of proteins on nascent DNA (aniPOND) is a powerful tool that has been used to identify viral and cellular proteins on nascent herpesvirus replication forks (32, 34, 35). Anipond analysis of HSV-1 infected cells identified host factors involved in BER, mismatch repair, and double-strand DNA break sensing along with viral replication factors on nascent viral DNA (32). KSHV LANA was observed to bind human UNG2 suggesting a requirement for uracil excision at origins of viral DNA replication (36). We previously reported that an MHV68 virus lacking the catalytic functions of both vUNG and vDUT exhibited defects in replication and latency establishment in infected mice (25). This double mutant also manifested a phenotype of genomic instability, revealed by rapid loss of a non-essential reporter gene upon serial passage in mice (25). Further studies of the association of vUNG with vPOL and vPPF in the context of uracilated DNA are needed to understand the role of vUNG in viral DNA replication and genomic integrity.

The vUNG catalytic mutant in our study contains mutations in both DNA binding and enzymatic domains, ablating catalytic activity in our previous report (25). These mutations did not impact the interaction or colocalization with vPOL and vPPF. Further, we observe no defect for replication in cell culture or pathogenesis in mice for the MHV68 vUNG.CM containing both H207L and D85N mutations, unless coupled with the dUTPase catalytic mutant (25). The H213L mutation in the putative DNA binding domain of EBV vUNG disrupts interaction with the EBV vPOL in the context of reactivation and this mutant exhibited a reactivation defect (26). A recent study with KSHV reported that reactivation of KSHV BAC DNA containing mutations in the putative DNA binding domain (H210L) of vUNG *(ORF46)*, but not the putative enzymatic domain (Q87L/D88N), resulted in a defect in viral DNA synthesis and virus production (37). A caveat of these previous studies is that the loss of biochemical functions were not validated. Future studies should carefully dissect the interaction between the vUNG DNA binding domain and uracilated viral genomes from its roles in uracil excision by pursuing cell culture models with skewed nucleotide pools reflective of restrictive cell types *in vivo*. Cells that overexpress the cellular APOBEC3s or AID cytidine deaminases may also recapitulate host intrinsic and immune defenses encountered in the host (12).

Overall, we report that the MHV68 vUNG has non-enzymatic functions to complex with its cognate vPOL and vPPF. Taking our findings for MHV68 vUNG into account with the broader herpesvirus field leads us to posit that the vUNG may coordinate with vPOL and vPPF to surveil and repair uracil residues in the viral genome. Since the vUNG is dispensable in many cell culture systems but not *in vivo*, interactions with the viral replication factors are likely critical in restrictive cell types with skewed nucleotide pools or in cells expressing APOBECs. In the case of a gammaherpesvirus infection of germinal center B cells, AID may serve as a source of cytidine deamination (38). Future studies that dissect residues of MHV68 vUNG required for biochemical functions from protein interactions will enable the role of complex formation with vPOL and vPPF to be tested in the MHV68 pathogenesis system.

## MATERIALS AND METHODS

### Cell culture

Immortalized murine fibroblast cells, NIH 3T3 and NIH 3T12, and HEK-293T cells were obtained from ATCC (Manassa, Virginia, USA) and maintained in Dulbecco’s modified Eagle’s medium (DMEM) supplemented with 8% (NIH 3T3 and NIH 3T12) or 10% (HEK-293T) fetal bovine serum, 1% L-glutamine, 100 U/ml penicillin, and 100 mg/ml streptomycin at 37°C in 5% CO_2_.

### Plasmid expression constructs

The FLAG-vPOL construct was generated by amplifying the MHV68 vPOL from the recombinant MHV68 M3-Luc bacterial artificial chromosome (BAC) (39) with PCR followed by insertion in the p3xFLAG-Myc-CMV-24 vector (CloneTech Takara Biotech, Mountainview, CA) at the EcoRI and SalI restriction sites. RFLP with EcoRV and XhoI was used to verify the inserts. The vPPF-V5 construct was a gift from Pinghui Feng Ph.D. (University of Southern California). The pEF1α-V5 empty vector plasmid was created by removal of the vPPF sequence from the vPPF-V5 construct at the KpnI and XhoI restriction sites, followed by an adaptor ligation reaction. RFLP with BamHI and XbaI was used to verify the inserts.

The vUNG-FLAG, vUNG-MYC, vUNG.CM-FLAG and vUNG.CM-MYC constructs were synthesized (Blue Heron Biotech, Bothell, WA). Briefly, the WT-vUNG (MHV68 *ORF46*) (GenBank accession no. CAA70280) and CM-vUNG genes were engineered with a C-terminal MYC or FLAG tag separated by a flexible GGSG amino acid linker and cloned into the OriGene pCMV6-AC expression vector. The vUNG catalytic mutant constructs were designed as previously described (25) with an aspartic acid in place of the wild-type asparagine (D85N) in the water activating loop, and a histidine (H207L) in place of the wild-type leucine in the DNA intercalating loop. The WT and CM vUNG constructs were validated by Oxford Nanopore plasmid sequencing.

### Recombinant viruses

The recombinant MHV68 H2b-YFP was previously described (40). The recombinant MHV68 with a catalytic mutation in the vUNG gene encoding vUNG (vUNG.CM) and the vUNG.CM marker rescue (MR) virus on the H2b-YFP reporter viral genome were previously described (25). The primers and gene blocks used in this study are listed in Table S1. The WT DsRed BAC and the BACs with a C-terminal GFP fusion to vPOL or vPPF were generated via BAC-mediated *en passant* mutagenesis (41). For the WT DsRed BAC, a targeting construct containing DsRed was designed to replace the GFP gene BAC cassette. Briefly, the DsRed gene was PCR amplifed from the pMSCV-IRES-DsRed Fp vector (Addgene, Watertown, MA, USA) and cloned via HindIII and KpnI into the pBlueScript KS+ vector to generate pBS_DsRed. A synthetic KanR cassette (gBLOCK) (42) (Integrated DNA Technologies, Coralville, IA) was constructed so that it contained the IscEI homing enzyme recognition site as well as a 50bp region homologous to the 50bp immediately downstream of the PstI restriction site of pBS_DsRed. The dsRed_KanR was PCR-amplified with primers designed with 20 bp homology to the flanking region of the desired insertion site (in this case, the GFP ORF in the BAC). The flanking regions up- and downstream of the GFP ORF were PCR-amplified with GFP_FLANK 1 and GFP_FLANK 2 forward and reverse primers. The PCR products were gel-purified (Qiagen) and assembled using Gibson Assembly (New England Biolabs). To generate the final targeting construct, a final PCR was performed using GFP_FLANK1_FOR and GFP_FLANK2_REV. This PCR product was gel-purified and electroporated into *E. coli* strain GS1783.5 cells harboring the WT MHV68 BAC. The KanR cassette was removed as outlined in the *en passant* protocol (41).

To generate vPOL-GFP and vPPF-GFP BACs, targeting constructs were designed to insert GFP as a fusion protein at the C-terminus of vPOL or vPPF. Two gBLOCKS containing a portion of EGFP and a kanamycin resistance cassette were synthesized and cloned into pBlueScript (KS+) to create the EGFP_KanR insert. Gene Block 1 was generated by PCR with Gene block 1 forward and reverse primers and the PCR product was digested with BamHI and HindIII and cloned into pBlueScript (KS+) to create pBS_Gene_block 1. Gene block 2 was generated by PCR with Gene block 2 forward and reverse primers and the PCR product was digested with HindIII and KpnI and cloned into pBS_Gene_block 1 to create pBS_Gene_block 2. pBS_Gene_block 2 was used as a template for PCR to amplify the EGFP_KanR cassette with forward and reverse primers for Gibson Assembly with regions consisting of approximately 600 bp flanking each insertion site of vPOL and vPPF. These regions were PCR amplified with vPOL_Flanking region 1 and 2 forward and reverse primers and vPPF_Flanking region 1 and 2 forward and reverse primers. The final targeting construct was generated by PCR using the outermost forward and reverse primers, gel-purified and electroporated into GS1783.5 harboring the MHV68 DsRed BAC.

Virus passage and titer determination were performed as previously described (43). RFLP with BamHI and HindIII, coupled with whole genome BAC sequencing (Illumina, San Diego, CA) verified that the insertion of vPOL-GFP and vPPF-GFP BACs did not lead to ORF coding changes or large-scale alterations of the viral genomes.

### Analysis of Recombinant Viral BAC DNA

To validate the sequences of recombinant viruses, BAC DNA was subjected to Illumina sequencing. Raw data, paired FASTQ files (SeqCenter, Pittsburgh, PA) was uploaded to the Galaxy server (usegalaxy.org). The Burrows-Wheeler Alignment tool: Map with BWA (Galaxy Version 0.7.17.5) was used with default settings to align sequencing reads against a reference MHV68 genome (U97553.2). A consensus sequence from the BAM output from the BWA algorithm was generated with the IVAR consensus tool (Galaxy Version 1.3.1+galaxy0). No SNPs were detected between the assembled and reference sequences by ClustalW (Galaxy Version 1.3.1+galaxy0) (44).

### Flow cytometric analysis of vUNG expression

NIH 3T12 cells were infected with recombinant MHV68 H2b-YFP at an MOI of 3. Cells were fixed and permeabilized before flow cytometry. Viability of the NIH 3T12 cells post infection, rate of infection and correlative degree of vUNG expression were recorded using the BD LSRFortessa (Franklin lakes, NJ, USA). Viability of NIH 3T12 cells was determined by propidium iodide negative cells. Rate of infection was determined by calculating the % of NIH-3T12 cells with YFP+ signal (Fig S1A). vUNG was detected by staining the cells with anti-vUNG-C1 and secondary anti-mouse AF647 (Jackson-Immuno, USA) prior to analysis. A minimum of two biological replicates were performed for each experimental analysis.

### Co-Immunoprecipitation and mass spectrometry following infection

NIH 3T12 cells were infected at an MOI of 3.0 and harvested at 36 hpi for co-IP/MS analysis. Cell pellets were washed with cold phosphate-buffered saline (PBS), followed by incubation with non-denaturing lysis buffer (20mM Tris HCl pH 8, 137mM NaCl, 10% glycerol, 1% NP-40, and 2mM EDTA) containing 50 μg/mL protease inhibitors (phenylmethylsulfonyl chloride), and 1 μg/mL aprotinin for 1 h at 4° C with constant agitation. The lysates were clarified by centrifugation, and the protein concentration of the supernatant was quantified by Bradford assay (ThermoFisher, Waltham, MA, USA). 4 μg of anti-vUNG-C1 or IgG isotype control was incubated with 4 mg of protein for 4 h at 4° C with gentle agitation prior to 1 h of incubation with 20 μL Praesto^®^ Jetted A50 protein A beads (Purolite, Montgomery, PA, USA), followed by washing with the non-denaturing lysis buffer three times. For elution, 50 μL of 2X SDS buffer was added to lysates prior to heating at 50° C for 10 min. The supernatant was collected and boiled with 100 mM DTT prior to SDS-PAGE to confirm vUNG capture by western or sent for mass spectrometry. Coomassie stained PAGE fragments were prepared as per instructions (UT Austin Center for Biomedical Research Support Biological Mass Spectrometry Facility, Austin, TX; RRID:SCR_021728).

Scaffold 5.0 (Proteome Software, Inc., Portland, USA) was used to probabilistically validate protein identifications derived from MS/MS sequencing results. Briefly, Scaffold verifies peptide IDs assigned by SEQUEST, Mascot, using the X!Tandem database (45, 46). The fold change or enrichment of peptides in the target sample (with αvUNG) was calculated as follows. First, for each peptide spectral match (PSMs), the false discovery rate (FDR) was set at 0.1% followed by a minimum cutoff of 10 unique peptides for any protein were considered for downstream analysis (Table S2). Further, with Scaffold 5.0, protein identifications were accepted if they could be established at greater than 99.0% probability of being protein and not a decoy. Second, non-specific binding of host protein from NIH 3T12 cells was addressed by discarding antigen IDs that were not enriched at least 3-fold in the infected sample when compared with the mock IP (Table S3). Finally, the Log_2_ fold change (FC) was calculated by comparing αvUNG and IgG isotype control (Table S4).

### Virus growth curve

NIH 3T12 cells were seeded at 1.8 x 10^5^ per well in 12-well plates 1 day prior to infection with recombinant MHV68 at an MOI of 3 in triplicate. Triplicate wells were harvested for each timepoint, and the cells and conditioned media were stored at 80°C and subjected to three cryogenic disruptions prior to the determination of titer by plaque assay (43).

### Antibodies and immunoblotting

For analysis of lytic antigen expression, infected cell pellets were lysed in RIPA buffer (ThermoFIsher) supplemented with protease inhibitors (Millipore Sigma). Protein concentration was quantified via BCA assay (Thermo Fisher) prior to SDS PAGE in 4-12% gradient gels (Invitrogen) with MOPs buffer (Invitrogen). Following electrophoresis, the gels were transferred to nitrocellulose membranes (Invitrogen) with an iBlot 2 dry blotting system (Thermo Scientific). Primary antibodies include vPPF, which was detected using an affinity-purified rabbit polyclonal anti-ORF59 antibody that was generated (GenScript, Piscataway, NJ, USA) against the peptides GKKTRGGNKASDSGT and KRPPPKKDREPTTKRPKL, corresponding to amino acids 210–224 and 372–388, respectively, of the predicted vPPF protein of MHV68 (Genbank:AAF19323) (47). vUNG was detected using anti-vUNG polyclonal sera (vUNG pAb) (25). GFP was detected with mouse anti-GFP mAb (Invitrogen, Clone GF28R, CA# MA5-15256), mouse sera harvested from MHV68-infected C57BL/6 mice 28 dpi was used to detect viral antigen. β-actin was detected with rabbit mAb (Cell Signaling, Danvers, MA, USA. Clone 13E5, CA# 4970). Secondary antibodies used were goat anti-mouse IgG HRP (Invitrogen, CA# 31430), goat anti-rabbit IgG HRP (Invitrogen, CA# 31480), and goat anti-rabbit IgG Alexa Fluor Plus 800 (Invitrogen, CA# A32735).

For analysis of Infected lysates (for input condition) and elutes from immunoprecipitation with protein A beads, samples were collected and 100 mM DTT was added. Samples were boiled and separated by 4-20% SDS PAGE prior to PVDF membrane transfer. Primary antibodies were the anti-vPPF pAb, the vUNG pAb, and mouse anti-tubulin mAb (clone 12G10, RAPC, MDACC). Secondary antibodies against mouse (Fcγ, cat # 115-005-164), rabbit (H+L, cat # 111-005-144), and human (H+L, cat # 109-005-088) were purchased from Jackson-Immuno Research and conjugated with Alexa Fluor 647 (Alexa Fluor™ Antibody Labeling Kits, cat # A20186). Since the anti-vUNG-C1 used for IP is on the mouse backbone, it was critical to avoid detection of light chain during immunoblotting of immunoprecipitated elutes since the light chain of C1 and vUNG has a similar electrophoretic mobility around 25-27KDa. Thus, we used anti-mouse Fcγ secondary antibody. A pilot antibody::protein A capture, as shown in Fig S1B, demonstrated the detection of only the heavy chain of anti-vUNG C1 immobilized by protein A (lane 3) from input fraction (lane 1). As such, during experimental detection of immunoprecipitated vUNG using mouse sera harvested from MHV68-infected C57BL/6 mice, the bands detected around 25-27KDa is vUNG and not the light chain.

For transfected lysates, protein was quantified via BCA assay (Thermo Fisher) and input and co-IP samples were subjected to SDS PAGE in 4-12% gradient gels (Invitrogen) with MOPs buffer (Invitrogen). Following electrophoresis, the gels were transferred to nitrocellulose membranes (Invitrogen) with an iBlot 2 dry blotting system (Thermo Scientific). Primary antibodies used were vUNG pAb (25), rabbit anti-FLAG mAb (Cell Signaling, Danvers, MA, USA; clone D6W5B, CA# 14793), mouse anti-V5 mAb (Biorad, Hercules, CA, USA; clone SV5-Pk1, CA# MCA1360), and rabbit anti-GAPDH mAb (Cell signaling; clone 14C10, CA# 2118). Secondary antibodies were goat anti-mouse IgG Alexa Fluor Plus 680 (Invitrogen, CA# A32729), and goat anti-rabbit IgG Alexa Fluor Plus 800 (Invitrogen, CA# A32735). Horseradish peroxidase-conjugated secondary antibodies were detected using the Pierce ECL Western Blotting Substrate (Thermo Scientific). Data were collected with the iBright™ FL1500 Imaging System (Invitrogen).

### Immunofluorescence microscopy

NIH 3T3 fibroblasts were seeded on glass coverslips (Marienfeld Superior, Lauda-Königshofen, Germany) in 24-well plates prior to infection at an MOI of 3. Cells were fixed in 4% paraformaldehyde in PBS for 10 min and permeabilized in 0.2% Triton X-100 in PBS for 5 min. Cells were blocked in 10% normal goat serum (Thermo Scientific, Waltham, MA) in PBS for 1 h, followed by incubation with primary antibodies diluted in blocking buffer for 1 h at room temperature with rocking, and then two PBS washes. Primary antibodies include anti-vPPF pAb, vUNG pAb, and anti-vUNG conjugated to Alexa-Fluor 647 (29). vPPF or vUNG was detected by incubation with the respective secondary antibodies, goat anti-mouse IgG conjugated to Alexa Fluor 568 (Invitrogen, CA# A11061) or goat anti-rabbit IgG Alexa Flour 555 (Invitrogen, CA# A27039) diluted in 10% blocking buffer for 1 h at room temp with rocking. Cells were stained with 4′,6′-diamidino-2-phenylindole (DAPI) (Thermo Fisher) nuclear stain diluted in PBS for 5 min with rocking. Coverslips were mounted using Prolong Gold antifade reagent (Invitrogen). Confocal microscopy was conducted using a spinning disk confocal with a Nikon T2i Eclipse Microscope (Nikon, Minato City, Tokyo, Japan). Z-stack images consisting of ten 2.5 μm optical sections were generated using Nikon NIS-Elements software and processed using ImageJ software (NIH, Bethesda, MD, USA). Line graphs of distance and intensity were generated using Nikon NIS-Elements software.

### Transfections

Plasmids were co-transfected into HEK-293T cells seeded at 7.5×10^5^ cells per well of a six-well dish in duplicate prior to transfection with TransIT-LT1 per manufacturer protocol (Mirus Bio, Madison, WI, USA). 24 h post-transfection duplicate wells were expanded into a 10 cm^2^ dish and incubated at 37°C for an additional 24 h.

### Co-immunoprecipitation

HEK-293Ts transfected with the indicated plasmids were lysed in M-PER lysis buffer (Thermo Scientific) supplemented with EDTA-free protease inhibitor (Roche, Indianapolis, IN) and 0.1 M NaCl. Lysates were sonicated with 5 pulses on ice using a Branson Sonifier 450 (duty cycle 10%, output control 2) and cleared by 14,000 x g centrifugation for 5 min at 4°C. Lysates were treated with 0.5 U DNase I per 100 uL lysate (Invitrogen, Waltham, MA, USA) for 15 min. For the analysis of vPPF interaction with vUNG, protein was lysed in M-PER buffer supplemented with EDTA-free protease inhibitor and 0.1 M NaCl and 0.1% Triton X-100. Lysates were treated with 10 U DNaseI per 100 uL lysate (Invitrogen) for 60 minutes at 37°C prior to IP steps.

For IP, lysates were incubated with primary antibodies (1:100) for 1 h with rotation at 4°C, and then rotated overnight at 4°C with 1.5 mg protein G (Dynabead, Invitrogen). Complexes were precipitated by centrifugation and washed three times in PBS supplemented 0.02% Tween 20 and resuspended in 100 uL PBS prior to elution with 40 uL of 2 parts 0.1M glycine pH 2.8 and 1 part NuPage sample buffer and reducing agent (Invitrogen). Samples were heated at 70°C for 10 min prior to magnetic removal of Dynabeads and storage at -20°C.

### Statistics

All data were analyzed using Prism 9 software (GraphPad, La Jolla, CA). Statistical significance was determined using two-way Anova with Tukey’s multiple comparisons test.

## ACKNOWLEDGEMENTS

We thank Dr. Michael Kruhlak of the CCR Confocal Microscopy Core facility and Dr. Jan Wizniewski of the NCI Experimental Immunology Branch Microscopy and Digital Imaging Facility for their assistance with confocal microscopy and image analysis. Thanks to Isabel Daher and Dr. Anna Grosskopf for technical assistance. This research was supported in part by the Intramural Research Programs of the NIH, the National Cancer Institute (NCI), Center for Cancer Research (K.R.S and L.T.K.). D.G.O. was supported by the Gundersen Medical Foundation. J.C.F. was supported by NIH/NCI R01CA167065. K.M.M. was supported by National Institutes of Health R01AI12539 and R21 Al111129.

K.R.S., S.P., Q.D., O.A., K.M.M., L.T.K., designed the experiments. K.R.S., S.P., O.A., D.G.O., executed the experiments. K.R.S., S.P., D.G.O., K.M.M., L.T.K., analyzed the data. K.R.S., S.P., K.M.M., L.T.K., prepared the manuscript.

## REFERENCES

1. Weller SK, Coen DM. 2012. Herpes simplex viruses: mechanisms of DNA replication. Cold Spring Harb Perspect Biol 4:a013011.

2. Wang Y, Tibbetts SA, Krug LT. 2021. Conquering the Host: Determinants of Pathogenesis Learned from Murine Gammaherpesvirus 68. Annual Review of Virology 8:349–371.

3. Pyles RB, Thompson RL. 1994. Mutations in accessory DNA replicating functions alter the relative mutation frequency of herpes simplex virus type 1 strains in cultured murine cells. J Virol 68:4514–24.

4. Prichard MN, Duke GM, Mocarski ES. 1996. Human cytomegalovirus uracil DNA glycosylase is required for the normal temporal regulation of both DNA synthesis and viral replication. J Virol 70:3018–25.

5. Courcelle CT, Courcelle J, Prichard MN, Mocarski ES. 2001. Requirement for uracil-DNA glycosylase during the transition to late-phase cytomegalovirus DNA replication. J Virol 75:7592–601.

6. Krokan HE, Bjørås M. 2013. Base excision repair. Cold Spring Harbor perspectives in biology 5:a012583.

7. Vértessy BG, Tóth J. 2009. Keeping uracil out of DNA: physiological role, structure and catalytic mechanism of dUTPases. Acc Chem Res 42:97–106.

8. Kavli B, Otterlei M, Slupphaug G, Krokan HE. 2007. Uracil in DNA—General mutagen, but normal intermediate in acquired immunity. DNA Repair 6:505–516.

9. Cheng AZ, Moraes SN, Shaban NM, Fanunza E, Bierle CJ, Southern PJ, Bresnahan WA, Rice SA, Harris RS. 2021. APOBECs and Herpesviruses. Viruses 13.

10. Suspène R, Aynaud M-M, Koch S, Pasdeloup D, Labetoulle M, Gaertner B, Vartanian J-P, Meyerhans A, Wain-Hobson S. 2011. Genetic Editing of Herpes Simplex Virus 1 and Epstein-Barr Herpesvirus Genomes by Human APOBEC3 Cytidine Deaminases in Culture and In Vivo. Journal of Virology 85:7594–7602.

11. Packard JE, Dembowski JA. 2021. HSV-1 DNA Replication—Coordinated Regulation by Viral and Cellular Factors. Viruses 13:2015.

12. Stewart JA, Damania B. 2023. Human DNA tumor viruses evade uracil-mediated antiviral immunity. PLoS pathogens 19:e1011252.

13. Cheng AZ, Yockteng-Melgar J, Jarvis MC, Malik-Soni N, Borozan I, Carpenter MA, McCann JL, Ebrahimi D, Shaban NM, Marcon E, Greenblatt J, Brown WL, Frappier L, Harris RS. 2019. Epstein-Barr virus BORF2 inhibits cellular APOBEC3B to preserve viral genome integrity. Nat Microbiol 4:78–88.

14. Kavli B, Sundheim O, Akbari M, Otterlei M, Nilsen H, Skorpen F, Aas PA, Hagen L, Krokan HE, Slupphaug G. 2002. hUNG2 is the major repair enzyme for removal of uracil from U:A matches, U:G mismatches, and U in single-stranded DNA, with hSMUG1 as a broad specificity backup. J Biol Chem 277:39926-36.

15. Zharkov DO, Mechetin GV, Nevinsky GA. 2010. Uracil-DNA glycosylase: Structural, thermodynamic and kinetic aspects of lesion search and recognition. Mutat Res 685:11–20.

16. Krokan HE, Drabløs F, Slupphaug G. 2002. Uracil in DNA – occurrence, consequences and repair. Oncogene 21:8935–8948.

17. Bogani F, Chua CN, Boehmer PE. 2009. Reconstitution of uracil DNA glycosylase-initiated base excision repair in herpes simplex virus-1. J Biol Chem 284:16784–16790.

18. Minkah N, Macaluso M, Oldenburg DG, Paden CR, White DW, McBride KM, Krug LT. 2015. Absence of the uracil DNA glycosylase of murine gammaherpesvirus 68 impairs replication and delays the establishment of latency in vivo. J Virol 89:3366–79.

19. Reddy SM, Williams M, Cohen JI. 1998. Expression of a uracil DNA glycosylase (UNG) inhibitor in mammalian cells: varicella-zoster virus can replicate in vitro in the absence of detectable UNG activity. Virology 251:393–401.

20. Ward TM, Williams MV, Traina-Dorge V, Gray WL. 2009. The simian varicella virus uracil DNA glycosylase and dUTPase genes are expressed in vivo, but are non-essential for replication in cell culture. Virus Research 142:78–84.

21. Mullaney J, Moss HW, McGeoch DJ. 1989. Gene UL2 of herpes simplex virus type 1 encodes a uracil-DNA glycosylase. J Gen Virol 70 (Pt 2):449–54.

22. Pyles RB, Thompson RL. 1994. Evidence that the herpes simplex virus type 1 uracil DNA glycosylase is required for efficient viral replication and latency in the murine nervous system. J Virol 68:4963–72.

23. Prichard MN, Lawlor H, Duke GM, Mo C, Wang Z, Dixon M, Kemble G, Kern ER. 2005. Human cytomegalovirus uracil DNA glycosylase associates with ppUL44 and accelerates the accumulation of viral DNA. Virol J 2:55.

24. Prichard MN, Duke GM, Mocarski ES. 1996. Human Cytomegalovirus Uracil DNA Glycosylase Is Required for the Normal Temporal Regulation of both DNA Synthesis and Viral Replication. Journal of Virology J Virol:3018–25.

25. Dong Q, Smith KR, Oldenburg DG, Shapiro M, Schutt WR, Malik L, Plummer JB, Mu Y, MacCarthy T, White DW, McBride KM, Krug LT. 2018. Combinatorial Loss of the Enzymatic Activities of Viral Uracil-DNA Glycosylase and Viral dUTPase Impairs Murine Gammaherpesvirus Pathogenesis and Leads to Increased Recombination-Based Deletion in the Viral Genome. mBio 9.

26. Su MT, Liu IH, Wu CW, Chang SM, Tsai CH, Yang PW, Chuang YC, Lee CP, Chen MR. 2014. Uracil DNA glycosylase BKRF3 contributes to Epstein-Barr virus DNA replication through physical interactions with proteins in viral DNA replication complex. J Virol 88:8883–99.

27. Bogani F, Corredeira I, Fernandez V, Sattler U, Rutvisuttinunt W, Defais M, Boehmer PE. 2010. Association between the herpes simplex virus-1 DNA polymerase and uracil DNA glycosylase. J Biol Chem 285:27664–72.

28. Strang BL, Coen DM. 2010. Interaction of the human cytomegalovirus uracil DNA glycosylase UL114 with the viral DNA polymerase catalytic subunit UL54. J Gen Virol 91:2029–2033.

29. Mu Y, Plummer JB, Zelazowska MA, Paul S, Dong Q, Chen Z, Krug LT, McBride KM. 2023. A Recombinant Antibody For Tracking Murine Gammaherpesvirus 68 Uracil DNA Glycosylase Expression. bioRxiv doi:10.1101/2023.05.17.541089:2023.05.17.541089.

30. Caragliano E, Bonazza S, Frascaroli G, Tang J, Soh TK, Grünewald K, Bosse JB, Brune W. 2022. Human cytomegalovirus forms phase-separated compartments at viral genomes to facilitate viral replication. Cell Reports 38:110469.

31. Ranneberg-Nilsen T, Dale HA, Luna L, Slettebakk R, Sundheim O, Rollag H, Bjoras M. 2008. Characterization of human cytomegalovirus uracil DNA glycosylase (UL114) and its interaction with polymerase processivity factor (UL44). J Mol Biol 381:276–88.

32. Dembowski JA, DeLuca NA. 2015. Selective Recruitment of Nuclear Factors to Productively Replicating Herpes Simplex Virus Genomes. PLOS Pathogens 11:e1004939.

33. Zeltzer S, Longmire P, Svoboda M, Bosco G, Goodrum F. 2022. Host translesion polymerases are required for viral genome integrity. Proc Natl Acad Sci U S A 119:e2203203119.

34. Dembowski JA, Dremel SE, DeLuca NA. 2017. Replication-Coupled Recruitment of Viral and Cellular Factors to Herpes Simplex Virus Type 1 Replication Forks for the Maintenance and Expression of Viral Genomes. PLoS Pathog 13:e1006166.

35. Manska S, Rossetto CC. 2022. Identification of cellular proteins associated with human cytomegalovirus (HCMV) DNA replication suggests novel cellular and viral interactions. Virology 566:26–41.

36. Verma SC, Bajaj BG, Cai Q, Si H, Seelhammer T, Robertson ES. 2006. Latency-associated nuclear antigen of Kaposi’s sarcoma-associated herpesvirus recruits uracil DNA glycosylase 2 at the terminal repeats and is important for latent persistence of the virus. J Virol 80:11178–90.

37. Morgens DW, Nandakumar D, Didychuk AL, Yang KJ, Glaunsinger BA. 2022. A Two-tiered functional screen identifies herpesviral transcriptional modifiers and their essential domains. PLoS Pathog 18:e1010236.

38. Bekerman E, Jeon D, Ardolino M, Coscoy L. 2013. A role for host activation-induced cytidine deaminase in innate immune defense against KSHV. PLoS pathogens 9:e1003748.

39. Hwang S, Wu TT, Tong LM, Kim KS, Martinez-Guzman D, Colantonio AD, Uittenbogaart CH, Sun R. 2008. Persistent gammaherpesvirus replication and dynamic interaction with the host in vivo. J Virol 82:12498–509.

40. Collins CM, Speck SH. 2012. Tracking murine gammaherpesvirus 68 infection of germinal center B cells in vivo. PLoS One 7:e33230.

41. Tischer BK, Smith GA, Osterrieder N. 2010. En passant mutagenesis: a two step markerless red recombination system. Methods Mol Biol 634:421–30.

42. Tischer BK, von Einem J, Kaufer B, Osterrieder N. 2006. Two-step red-mediated recombination for versatile high-efficiency markerless DNA manipulation in Escherichia coli. Biotechniques 40:191–7.

43. Cieniewicz B, Kirillov V, Daher I, Li X, Oldenburg Darby G, Dong Q, Bettke Julie A, Marcu Kenneth B, Krug Laurie T. 2022. IKKα-Mediated Noncanonical NF-κB Signaling Is Required To Support Murine Gammaherpesvirus 68 Latency In Vivo. Journal of Virology 96:e00027–22.

44. The Galaxy C. 2022. The Galaxy platform for accessible, reproducible and collaborative biomedical analyses: 2022 update. Nucleic Acids Research 50:W345–W351.

45. Craig R, Beavis RC. 2003. A method for reducing the time required to match protein sequences with tandem mass spectra. Rapid Commun Mass Spectrom 17:2310–6.

46. Searle BC. 2010. Scaffold: a bioinformatic tool for validating MS/MS-based proteomic studies. Proteomics 10:1265–9.

47. Upton JW, van Dyk LF, Speck SH. 2005. Characterization of murine gammaherpesvirus 68 v-cyclin interactions with cellular cdks. Virology 341:271–283.

